# Molecular and cellular correlates of human nerve regeneration: *ADCYAP1* encoding PACAP enhances sensory neuron outgrowth

**DOI:** 10.1101/686618

**Authors:** Annina B Schmid, Georgios Baskozos, Katherine Windsor, Pall Karlsson, Oliver Sandy-Hindmarch, Greg A Weir, Lucy A McDermott, Alex J Clark, Joanna Burchall, Akira Wiberg, Dominic Furniss, David LH Bennett

## Abstract

We only have a rudimentary understanding of the molecular and cellular determinants of human nerve regeneration. Here, we use carpal tunnel syndrome (CTS) as a human model system to prospectively evaluate correlates of neural regeneration and their relationship with clinical recovery after decompression surgery. At 6 months post-surgery, we noted a significant improvement of median nerve neurophysiological and somatosensory function. Serial skin biopsies revealed a partial recovery of intraepidermal innervation, whose extent correlated with symptom improvement. In myelinated afferents, nodal length increased postoperatively. Transcriptional profiling of the skin revealed 23 differentially expressed genes following decompression, with *ADCYAP1* (encoding PACAP) being the most strongly upregulated and showing an association with regeneration of intraepidermal nerve fibres. Using human induced pluripotent stem cell-derived sensory neurons, we confirmed that PACAP significantly enhances axon outgrowth *in vivo*. Since PACAP signals through G-protein receptors, this pathway provides an interesting therapeutic target for human sensory nerve regeneration.

## Introduction

Knowledge of cellular and molecular correlates of nerve regeneration is critical to understand the temporal profile of nerve regeneration as well as the determinants of successful recovery. While neural regeneration has been extensively studied in animal models of peripheral nerve injury^1, 2^, studying the determinants of nerve regeneration and its relationship to recovery in humans remains challenging due to the limited access to tissues and a lack of treatments to promote nerve regeneration.

So far, studies have mostly relied on sensory or motor reinnervation times, the advancement of Tinel’s sign as well as the recovery of action potentials in electrophysiological recordings^3^. In the past decade, serial skin biopsies have provided insights into the regeneration of small sensory axons in humans^4, 5, 6, 7^. Recently, skin biopsies have also been used to obtain information about the integrity of myelination and large nerve fibres^8, 9^. However, a lack of knowledge remains concerning the molecular and cellular determinants of nerve regeneration and their relationship to clinical recovery and specifically neuropathic pain in humans.

Here, we use carpal tunnel syndrome (CTS) as a model system to study sensory nerve regeneration in humans. CTS is one of the few neuropathic conditions that can be treated surgically. It therefore provides a unique model system to prospectively study the cellular and molecular correlates of nerve regeneration and their relationship to clinical recovery in humans. We deeply phenotyped patients before and six months after carpal tunnel decompression including electrophysiology, quantitative sensory testing, symptom profile and functional deficits. Using serial skin biopsies, we determined the postoperative reinnervation of sensory target tissue with histology as well as the molecular signature associated with regeneration using RNA-sequencing. This revealed that *ADCYAP1* was the most differentially expressed gene and the protein this gene encodes (PACAP) was found to enhance neurite outgrowth in human induced pluripotent stem cell-derived sensory neurones.

## Results

### The majority of patients show significant symptomatic and functional improvement

Sixty patients with electrodiagnostically confirmed CTS (mean age 62.5 (SD 12.2), 36 females) were examined before and six months after decompression surgery. Twenty healthy volunteers served as healthy controls. Baseline demographic and clinical data are presented in Table 1. Post-surgically, 83.3% of patients rated their symptoms at least ‘a good deal better’ (≥5) on the global rating of change scale. None of the patients experienced a deterioration of symptoms. In relation to hand function, 63.3% of operated patients felt at least ‘a good deal better’. The self-perceived symptoms and function improvements were consistent with a post-operative decrease of the Boston carpal tunnel questionnaire symptom and function subscales (Boston symptoms mean (SD) pre 2.8 (0.7), post 1.5 (0.5), t (59)=13.8, p<0.0001; Boston function pre 2.2 (0.8), post 1.5 (0.5), t (59)=7.00, p<0.0001). However, symptoms did not fully resolve in all patients, with 46.6% continuing to have some pain and 38.3% to feel paraesthesia or numbness, although these residual symptoms were mostly mild in nature.

**Table 1:** Demographic and clinical data of patients with carpal tunnel syndrome and healthy volunteers.

|  | surgery | no-surgery | controls | p-value |
| --- | --- | --- | --- | --- |
| Number of participants | 60 | 13 | 20 |  |
| Gender (female/male) | 36/24<br>(60.0% ♀) | 9/4<br>(69.2% ♀) | 14/6<br>(70.0% ♀) | 0.649□ |
| Mean Age (SD) [years] | 62.5 (12.2) | 53.6 (11.9)* | 58.0 (12.9) | 0.044 |
| Mean weight (SD) [kg] | 74.4 (15.0) | 74.1 (13.1) | 73.6 (15.7) | 0.978 |
| Mean height (SD) [cm] | 168.4 (9.5) | 165.2 (8.8) | 167.0 (7.6) | 0.472 |
| Mean symptom duration (SD)<br>[months] | 64.2 (96.5) | 52.7 (44.8) |  | 0.676 |
| Mean Boston scale (SD) |  |  |  |  |
| Symptoms | 2.8 (0.7) | 2.3 (0.6)* |  | 0.008 |
| Function | 2.2 (0.8) | 1.9 (0.6) |  | 0.184 |

SD: standard deviation; NPSI: neuropathic pain symptom inventory
\* Indicates statistical significance compared to surgery group.
Comparisons are done with one way ANOVA and independent t-tests except for □ which used chi-squared test.

### Electrodiagnostic test severity improves following surgery, but fails to reach normal levels in most patients

Electrodiagnostic testing demonstrated moderate severity preoperatively (median Bland scale 3.0 [IQR 2.8]). After surgery, electrodiagnostic test severity improved by one grade to an average of mild severity (2.0 [2.0], z(55)=-5.62, p<0.0001, Wilcoxon signed rank test, Table 2). With the exception of eight patients (4.8%), in whom electrodiagnostic testing was within normal limits following surgery, all patients continued to show some degree of median nerve conduction slowing.

**Table 2:**
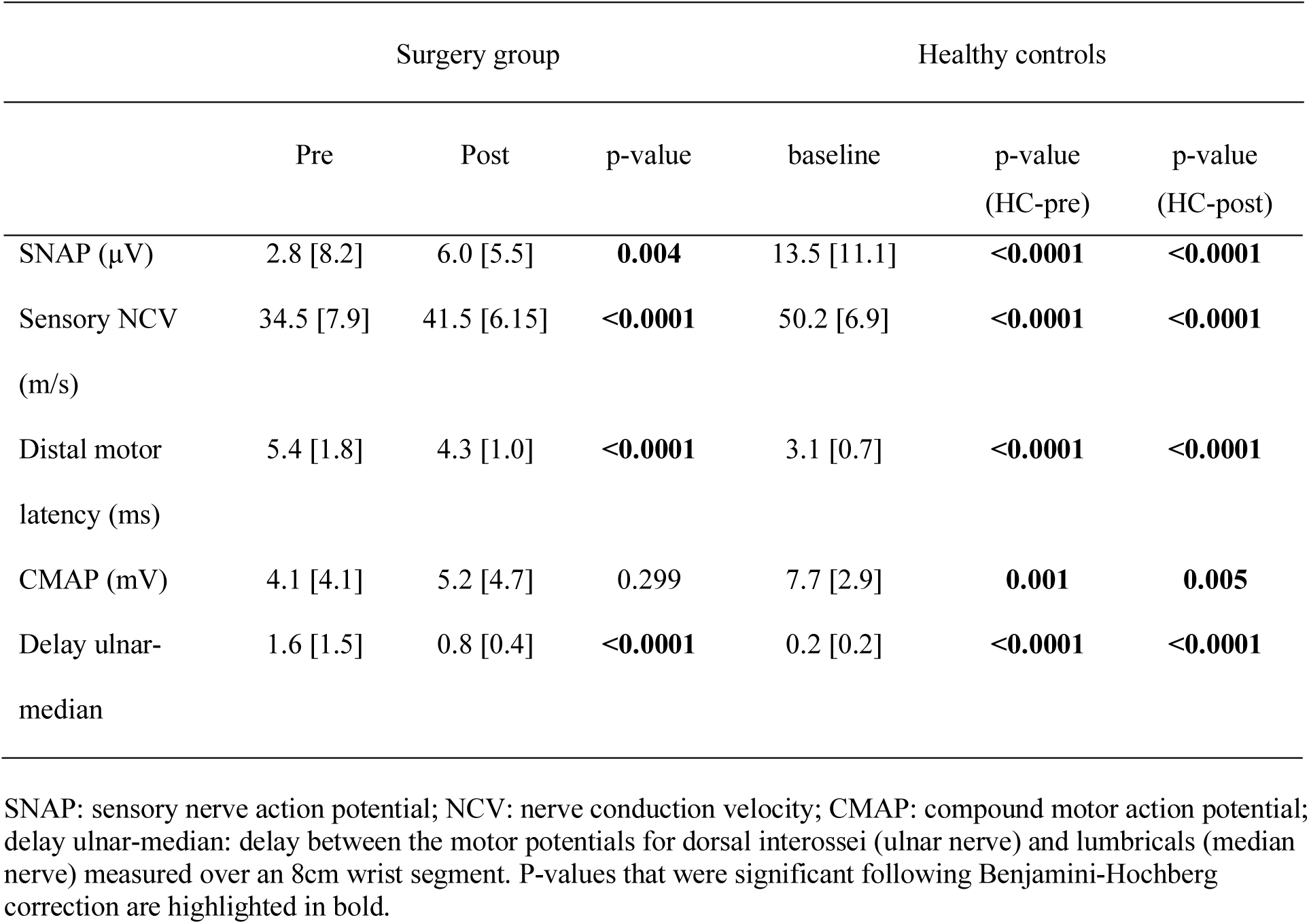
Median nerve neurophysiology data of patients with CTS pre- and post-surgery. Data are presented as median [interquartile range].

### Sensory detection thresholds improve following surgery, but fail to reach normal levels

Quantitative sensory testing data from the median innervated territory of the hand can be found in Table S1 and Fig 1A-C. Patients with CTS had a significant pre-surgical deficit in all thermal and mechanical detection thresholds compared to healthy controls (t(78)>2.30, p<0.015). Cold detection, thermal sensory limen, mechanical and vibration detection improved following surgery (t(59)>3.20, p<0.002). Only warm detection thresholds remained unchanged post-surgically (t(59)=1.58, p=0.120). Despite the improvement in most parameters, postoperative detection thresholds failed to reach levels found in healthy control participants (t(75)>2.00m p<0.033), except for vibration detection thresholds (t(74)=1.09, p=0.429).

**Fig 1:**
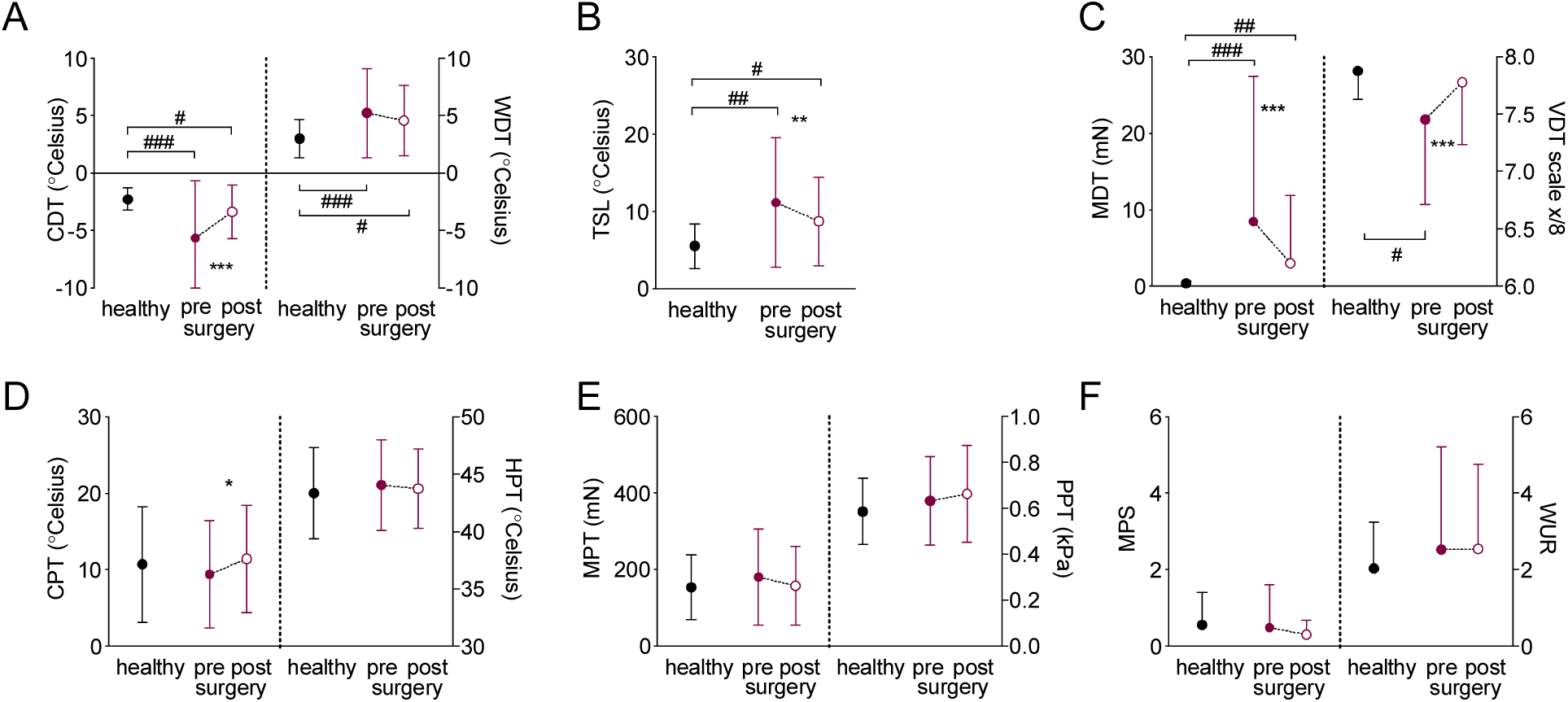
Recovery of somatosensory function. Raw quantitative sensory testing data are presented as mean and standard deviations for healthy participants (n=20, black circles) and patients with CTS before (closed red circle) and after surgery (n=60, open red circle). **(A)** thermal detection thresholds (cold (CDT) and warm detection thresholds (WDT)), **(B)** thermal sensory limen (TSL), **(C)** mechanical detection thresholds (mechanical (MDT) and vibration detection thresholds (VDT)), **(D)** thermal pain thresholds (cold (CPT) and heat pain thresholds (HPT)), **(E)** mechanical pain thresholds (mechanical (MPT) and pressure pain thresholds (PPT), **(F)** mechanical pain sensitivity (MPS) and windup ratio (WUR). Significant differences for paired/independent t-tests after Benjamini Hochberg correction are indicated between groups with #p<0.05 ##p<0.01 ###p<0.0001 and within groups with *p<0.05 **p<0.01 ***p<0.0001.

Thermal and mechanical pain thresholds were not different at group level between healthy controls and preoperative CTS patients (p>0.402, Fig 1D-F) and remained largely unaltered following surgery (p>0.221). The only exception was an increased sensitivity to cold pain following surgery compared to before surgery (t(59)=-2.04, p=0.046). Out of four patients with preoperative paradoxical heat sensations, one patient continued to paradoxically feel heat on cooling postoperatively and two patients experienced new paradoxical heat sensations postoperatively. No patient presented with dynamic mechanical allodynia either pre- or postoperatively.

### Intraepidermal nerve fibres partially regenerate following surgery

Skin biopsies in the median nerve innervated territory of the hand showed a substantial structural degeneration of small axons in the epidermal layer of patients’ skin compared to healthy controls (fibres/mm epidermis in CTS pre-operative mean (SD) 4.20 (2.83); healthy controls 8.03 (2.08), t(77)=5.3, p<0.0001, Figure 2A-C). Following surgery, intraepidermal nerve fibre density improved (5.35 (3.34), t(57)=-3.5, p=0.001), but failed to reach normal innervation levels (t(77)=3.4, p=0.001). Of note, postoperative regeneration was highly variable (Fig 2D).

**Fig 2:**
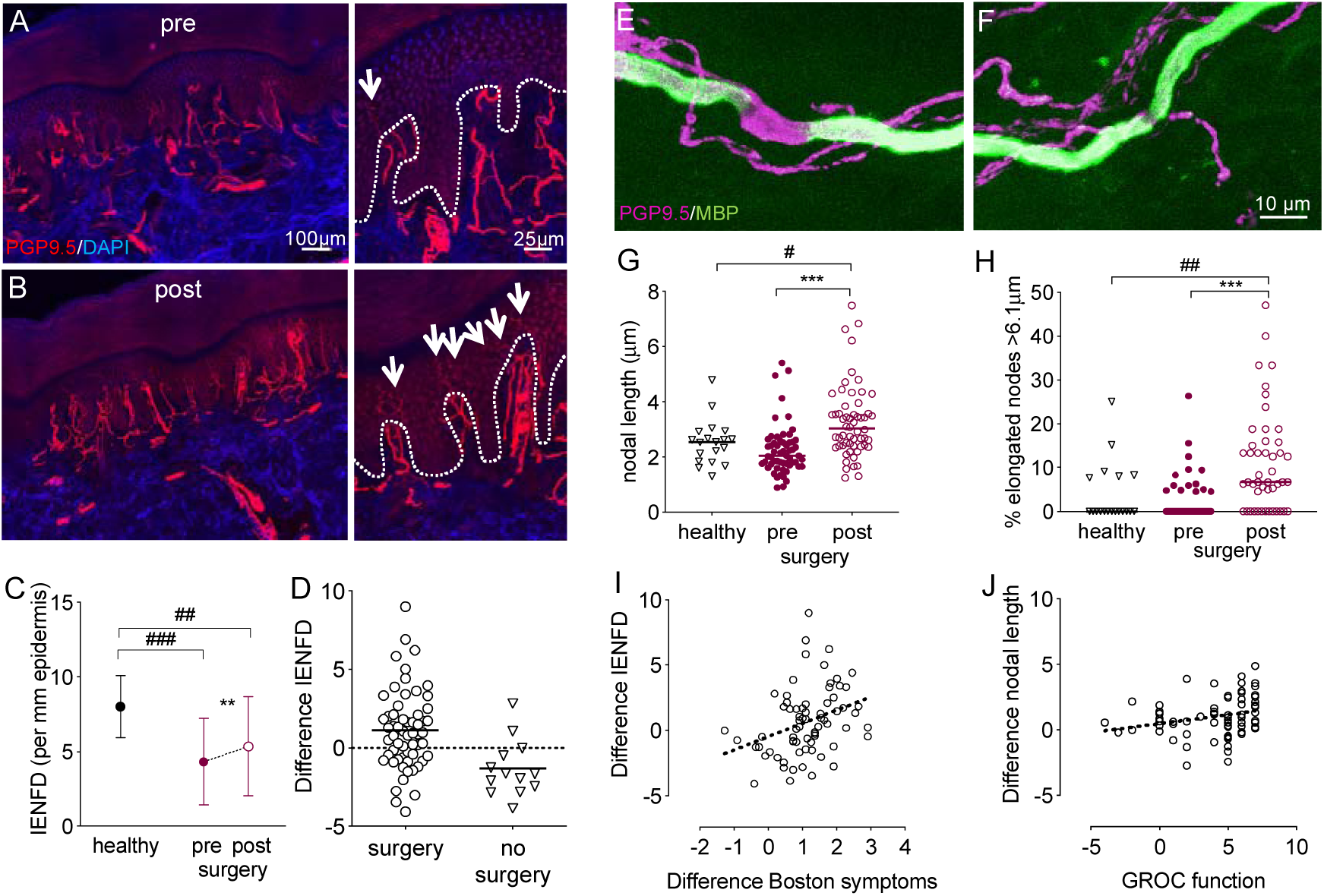
Histological evidence for neural regeneration into target tissue. A-D: small fibre regeneration. Immunohistochemically stained sections of index finger skin of a patient with CTS before **(A)** and 6 months after surgery **(B)**. Cell nuclei are apparent in blue (DAPI) and axons in red (PGP9.5). After surgery, an increased number of small fibres (arrows on 4x magnified regions of interest on the right) penetrate the dermal-epidermal border (dotted line). **(C)** quantification revealed a reduced intraepidermal nerve fibre density (IENFD) in patients with CTS compared to healthy controls, which increased after surgery but failed to reach normal levels. **(D)** The difference in IENFD post compared to pre surgery demonstrated a high interindividual variability, with a continuing small fibre degeneration in patients who were not operated. (E-H) Changes in nodal architecture. Example of an elongated **(E)** and normal node **(F)** in skin sections stained immunohistochemically with protein gene product 9.5 (PGP9.5) and myelin basic protein (MBP). **(G)** Median nodal length and **(H)** the median percentage of elongated nodes >6.1 μm increase following surgery and remain higher than healthy controls. Significant differences for paired/independent t-tests after Benjamini Hochberg correction are indicated between groups with #p<0.05 ##p<0.01 ###p<0.0001 and within groups with **p<0.01 ***p<0.0001. Group differences based on n=59 patients with CTS and n=20 healthy controls. **(I)** A more pronounced intraepidermal nerve fibre regeneration (Difference IENFD) correlates with improved symptoms (r=0.389, p=0.001, difference Boston symptom scale data multiplied with -1). **(J)** The correlation between difference in mean nodal length and the global rating of change scale (GROC) for hand function marginally fails to reach significance after Benjamini Hochberg correction (r=0.316, p=0.008).

### Subepidermal plexus nerve fibre length density remains unchanged

Given the incomplete epidermal reinnervation following surgery, we hypothesised that regenerating axons may reach the dermis but their penetration through the dermal-epidermal junction may be delayed^7, 10^. However, evaluation of nerve fibre length density in the subepidermal plexus did not reveal any differences in patients pre- and post-surgery (median [IQR] pre 70.25 [48.98]; post 75.85 [63.95], z(57)=1.07, p=0.285, Fig S1A).

### Nodal architecture changes following surgery

To evaluate changes in myelinated fibres, we examined nodal architecture which was previously found altered in patients with CTS (*9*) (Fig 2E-F). At baseline, the median nodal length (healthy 2.54 IQR [0.98]; CTS 2.03 [0.82], z(77)=-1.70, p=0.088) and the percentage of elongated nodes >6.1μm (healthy 0.00 [7.92]; CTS 0.00 [4.76], z(66)=-0.30, p=0.767) were comparable in patients with CTS and healthy controls. Following surgery, median nodal length (3.03 [1.23], z(56)=4.36, z(56)=4.36, p<0.0001) and the percentage of elongated nodes increased in patients (post 6.66 [15.79], z(46)=4.10, p<0.0001) and remained higher than healthy controls (z(77)=2.58, p=0.010 and z(66)=2.85, p=0.004 respectively, Fig 2G-H).

The density of Meissner corpuscles, the number of PGP+ axon bundles per mm^2^ dermis and the ratio of dermal PGP+ axon bundles containing myelin remained unchanged at baseline and after surgery (Fig S1B-D).

### Deterioration in small fibre integrity in patients not undergoing surgery

A small number of patients (n=13) opted not to undergo surgery. These were slightly younger and had less severe symptoms at baseline compared to those undergoing surgery (Table 1). Of note, non-operated patients demonstrated a continuing degeneration of their IENFD over time (mean baseline 6.94 (SD 3.09); follow up 5.61 (3.41), t(12)=2.66, p=0.021, Fig 2D). Data for the other clinical and histological measures of non-operated patients are summarized in Table S2.

### Neural regeneration correlates with symptom improvement

The extent of small fibre regeneration positively correlates with symptom improvement as determined with the Boston symptom questionnaire (r=0.389, p=0.001, Fig 2I). A postoperative increase in nodal length (r=0.302, p=0.012, Fig 2J) and percentage of elongated nodes with GROC for hand function marginally failed to reach significance after Benjamini Hochberg correction (r=0.306, p=0.019). None of the other correlations were significant (p>0.144).

### 31 genes including ADCYAP1 are differentially expressed following surgery

To determine a molecular signature associated with nerve regeneration, we performed RNA sequencing of the skin of 47 patients with CTS (n=29 females) before and after surgical decompression. Thirty-one genes were significantly differentially expressed (DE) in CTS patients post-versus pre-surgery (Fig 3A-B, Fig S2, Table S3). Gene ontology enrichment analysis for biological processes showed that these genes were enriched amongst others in proximal/distal pattern formation (Fig 3C, Table S4). The most significantly dysregulated gene was *ADCYAP1* (log2 fold change =1.87, FDR p-value = 0.0001, Fig 3A). *ADCYAP1* encodes PACAP (pituitary adenylate cyclase-activating peptide), which is a highly evolutionary conserved protein that is involved in a range of physiological processes including neuronal survival after injury and neurite outgrowth^11^. We confirmed the increased expression of *ADCYAP1* in skin samples following surgical release using the independent technique of droplet digital PCR (Fig S3).

**Fig 3:**
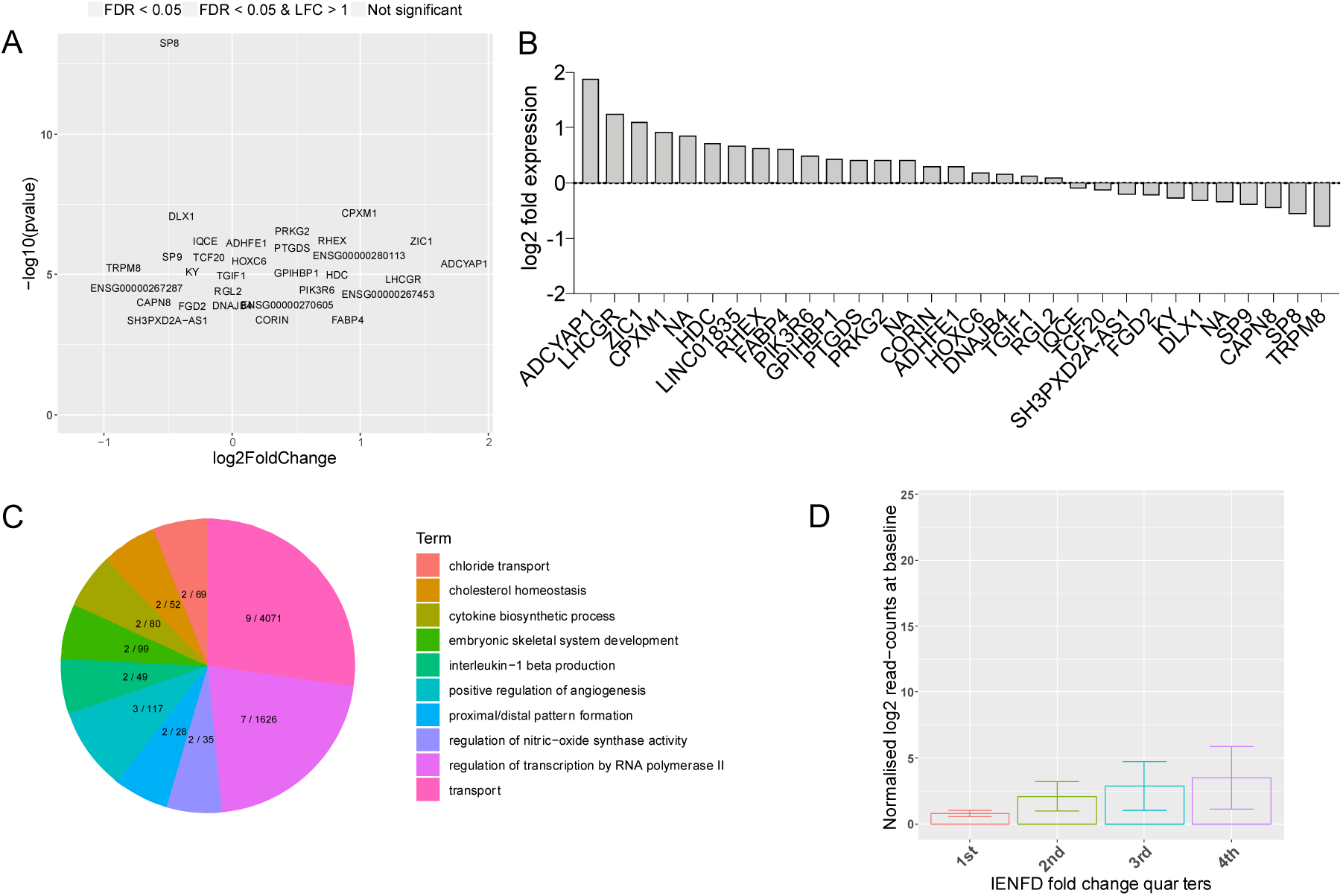
Molecular signature for nerve regeneration in skin. **(A)** Volcano plot showing the range of transcriptional changes following carpal tunnel surgery compared to before surgery. X-axis Log2 (Fold Change), y-axis -log10 (p-value). Significantly DE genes according to FDR adjusted p.value are highlighted and labelled. *ADCYAP1*, i.e. the gene encoding PACAP, was the most up-regulated gene. **(B)** Gene expression changes (log2 fold change) for DE expressed genes. **(C)** Top 20 biological process gene ontology (GO) terms from GO enrichment analysis with number of DE genes of total genes marked for each term on the pie chart. **(D)** *ADCYAP1* is upregulated at baseline in all IENFD fold change quarters compared to the lowest quarter of patients with negative IENFD fold change, who continue to degenerate after surgery. Graph shows mean normalised log2 read counts at baseline, standard errors and single data po

### Expression of DE genes is associated with histological evidence for nerve regeneration

We did not identify FDR adjusted significant correlations on the individual level between the RNA-sequencing-determined fold change of the 31 significantly DE genes and the 1) fold change or 2) differences in IENFD and mean nodal length post-surgery versus pre-surgery. We also defined four phenotypic groups (quarters) based on the continuous IENFD and nodal length fold change quartiles and tested for significant associations of these groups with baseline DE gene expression. Whereas no relationship was detected for nodal length, this analysis revealed that IENFD fold change quarters were significantly dependent on *ADCYAP1, DLX1, PRKG2, ADHFE1, and PIK361* baseline expression (p<0.05). *ADCYAP1* (Fig 3D) and *ADHFE1* are upregulated in all IENFD fold change quarters compared to the lowest quarter of patients with negative IENFD fold change, i.e. those who continue to degenerate. *PRKG2* is upregulated and *DLX1* and *PIK3R6* are downregulated in patients belonging to the 3^rd^ and 4^th^ quarter.

### PACAP is expressed in human skin and facilitates neurite outgrowth in vitro

We confirmed the localisation of PACAP within sensory afferents in human skin using immunohistochemistry (Fig 4A). Human induced pluripotent stem cell-derived (hiPSCd) sensory neurons have been shown to replicate many of the molecular, morphological and functional features of native human sensory neurons^12, 13, 14^ and so we used these to study the functional effects of PACAP. These hiPSCd-sensory neurons demonstrated strong mRNA expression of both *ADCYAP1*, as well as its high affinity receptor *PAC1* (Fig 4B). mRNA of the secondary receptors *VPAC1* and *VPAC2* with affinity for PACAP and VIP (vasoactive intestinal protein) are also expressed in hiPSCd-sensory neurons albeit at lower levels. We therefore used these neurons as an experimental *in vitro* model to explore a potential role of PACAP in neurite regeneration following axotomy. Immunostaining for PAC1 revealed that this was predominantly located in cell bodies and growth cones (Fig 4C), consistent with a role in promoting axon outgrowth. Application of 10 nM PACAP to hiPSCd-sensory neurons increased total neurite outgrowth length by 86% compared to vehicle (mean (SD) vehicle 570.9μm (181.8); PACAP 1065.0µm (285.5), t(11)=3.79, p=0.003, Fig 4D-E), confirming a pro-regenerative role of PACAP in human sensory neurons.

**Fig 4:**
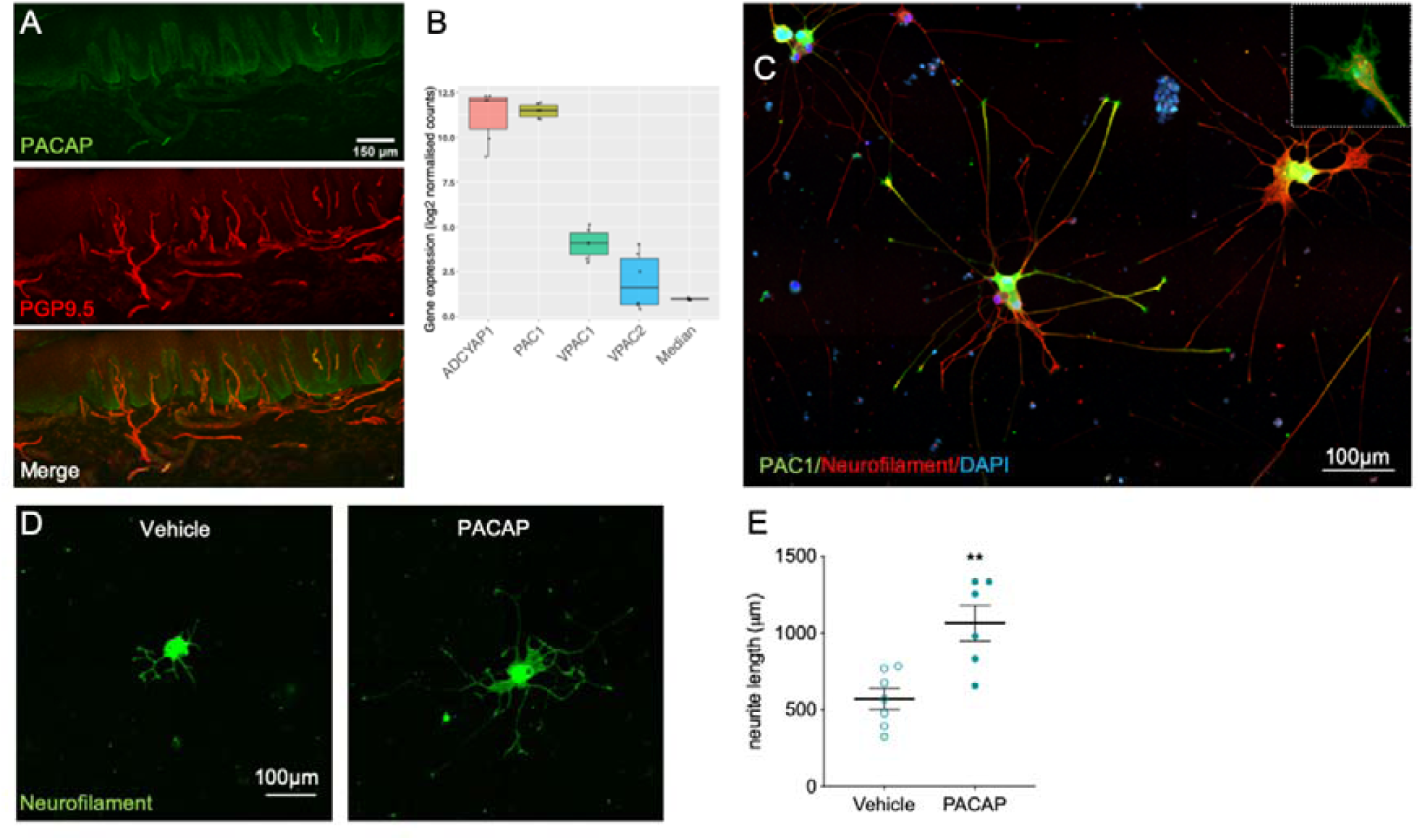
PACAP induces neurite outgrowth in human induced pluripotent stem-cell-derived (hiPSCd) sensory neurons. **(A)** Immunohistochemical staining in human finger skin demonstrating presence of PACAP (green) within sensory nerve fibres (PGP9.5). **(B)** Expression strength of *ADCYAP1*, its main receptor *PAC1* as well as secondary receptors *VPAC1 and VPAC2* and the median expression of all genes in hiPSCd-sensory neurones. Y-axis shows RNA-seq log2 normalised gene counts. Boxplots show the median and interquantile range with single data points depicting each experiment (n=3 cell lines, 2 differentiation each). **(C)** Immunohistochemical staining of hiPSCd-sensory neurons demonstrating expression of the PACAP receptor (PAC1, green) in the soma as well as at the growth cones (5x magnified inset). **(D)** PACAP given to dissociated mature and re-plated hiPSCd-sensory neurons enhances neurite outgrowth compared to vehicle. Data is quantified with mean and standard error of measurements in **(F)**, **=p=0.008

## Discussion

Little is known about the molecular and cellular determinants of nerve regeneration in humans. Using CTS as a model system, we demonstrate recovery of large and small fibre function following surgical nerve decompression; however, this recovery remains incomplete even 6 months after surgery. There was significant individual variation in the degree of cutaneous reinnervation, and at group level this did not reach the level seen in unaffected control individuals. We identified a significant correlation between both cutaneous reinnervation by small fibres and the nodal length of myelinated sensory fibres, with symptom and functional recovery following carpal tunnel decompression, respectively. Gene expression analysis identified 31 differentially expressed genes, with *ADCYAP1* encoding PACAP being most strongly upregulated. Intriguingly, *ADCYAP1* expression was associated with a histological regenerative phenotype and a regenerative role of PACAP was confirmed in hiPSCd-sensory neurons.

Carpal tunnel surgery greatly improved symptoms in the majority of patients (83%), which is in line with previous reports of excellent results in ∼75% of patients^15^. The subjective improvement was accompanied by a significant improvement in somatosensory function for all detection thresholds except warm detection. A selectively impaired postoperative recovery of warm detection has previously been reported in patients with entrapment neuropathies including CTS^16^ and lumbar radiculopathy^17^. Whereas this may be attributed to more subtle deficits in warm than cold detection in patients with CTS^18^, it may also be explained by a previously observed delayed functional recovery of unmyelinated C-fibres (encoding warm sensations) compared to A-fibres^19, 20^.

At 6 months after surgery, there was a significant reinnervation of the skin by epidermal nerve fibres, however this remained incomplete. Given the relatively short reinnervation distance and the previously reported rate of axon regeneration^21^ one may have expected a higher rate of reinnervation. However, this slow regeneration of small epidermal fibres is in line with previous animal work^22, 23^ and experimental human studies^4, 5^. It has been suggested that the epidermal layer of the skin may be a relatively hostile environment for regenerating fibres as demonstrated by incomplete epidermal regeneration while dermal reinnervation is successful^7, 10^. Here, we could not find a postoperative increase in the subepidermal plexus length or number of dermal axon bundles. The available quantification methods may however not be sensitive to detect relatively subtle changes in dermal small fibres amidst the abundance of large fibres.

Alternatively, dermal reinnervation may indeed remain incomplete or only reach subdermal levels, as previously shown for SP+ axons (small peptidergic fibres) following experimental nerve injury^23^. The slow rate of reinnervation emphasizes the point that any future trials of agents to promote nerve regeneration in which epidermal nerve fibres are used as an outcome measure would need to be of significant duration.

Meissner corpuscle and dermal myelinated bundle density in CTS patients were comparable to controls at baseline and remained unaltered after surgery despite a clear improvement in large fibre function assessed by QST (mechanical and vibration detection thresholds). Presumably, the large fibre dysfunction in CTS is driven by ischaemic and demyelinating changes at the lesion site - as has been shown in experimental^24^ and clinical nerve compression^25^ - rather than axon degeneration and changes to target tissue receptors^9, 24^. This is supported by the significant albeit incomplete recovery of electrophysiological properties over the affected wrist segment following surgical decompression. Persistence of thin myelin sheets after remyelination as shown in experimental models^26, 27^ may account for the continuing conduction slowing.

Unlike in a previous cohort^9^, patients with CTS did not have an increased nodal length at baseline. Since the presence of elongated nodes seems to be protective in nature^9^, this disparity may be caused by the inclusion of patients with more severe symptoms in the current pre-surgical cohort (not showing protective nodal elongation) compared to the patients with milder symptomatology in the previous cohort^9^. Of note, nodal length significantly increased following surgery, further corroborating a protective effect. The mechanism underlying the presence of elongated nodes remains speculative, but it could be associated with the dynamic process of demyelination or remyelination^28^. Given the increased presence of elongated nodes after surgery and their correlation with functional recovery, an association with remyelination seems more likely. Alternatively, it has been demonstrated that the incorporation of cytoskeletal and ion channel components to nodes of Ranvier requires vesicular transport^29^. It is therefore possible that the restoration of axon transport following carpal tunnel surgery enhances delivery of nodal components and nodal lengthening. Elongated nodes may also reflect nerve fibre lengthening during nerve regeneration, as has been shown in experimental limb lengthening in the rabbit^30^. The axon and its associated Schwann cell are closely linked at the septate-like junction of the paranode. Lengthening may therefore be more easily achieved at the node rather than at the internode, which would require increased length of both axon and Schwann cell.

We identified a large degree of interindividual variation in the extent of small fibre regeneration and nodal changes, which correlate with clinical recovery. Currently, therapeutic targets that promote nerve regeneration in humans are lacking. Using CTS as a model system, we hypothesised that we could identify a transcriptional signature in the skin that would show a relationship to successful reinnervation by sensory axons. We identified *ADCYAP1* as the most upregulated gene (3.7 fold change) using RNA-seq, and droplet digital PCR confirmed this up-regulation. This increased expression was associated with histological evidence of neural regeneration. *ADCYAP1* encodes PACAP (pituitary adenylate cyclase-activating peptide), which is a pleiotropic secreted molecule that has been associated with cell survival in neurodegenerative conditions^31^ as well as growth cone guidance^32^ and increased neurite outgrowth of peripheral afferent neurons in preclinical models^32, 33, 34, 35^. Animal studies have shown that PACAP is expressed in small to medium sized DRG neurons and upregulated during the regenerative process following nerve injury^36, 37^. In relation to localization, we could identify PACAP in sensory afferents in human skin. In terms of the source of this PACAP, it may be expressed by keratinocytes, released and subsequently bind to its receptor PAC1 (which we found was expressed on the growth cones of hiPSCd-sensory neurons). Alternatively, we also found that *ADCYAP1* mRNA is expressed by hiPSCd-sensory neurons and so it may be having an autocrine action on these neurons.

Whereas a regenerative function of PACAP on sensory neurons has been demonstrated for central^38, 39^ and peripheral neurons^33, 34, 35, 40^ in animals, its effects on neurite outgrowth in human neurons has not previously been tested. One study reported that PACAP can rescue hiPSCd-motor neurons from apoptosis^31^. We found that administration of PACAP to hiPSCd-sensory neurons resulted in a large (70%) and significant increase in neurite outgrowth *in vitro*. It should be noted that we found a number of other genes significantly associated with cutaneous reinnervation (e.g., *DLX1, PRKG2, ADHFE1, LINC01835, PIK361*) and this dataset can therefore be further exploited to identify biomarkers of and therapeutic targets relating to neural regeneration in humans.

Due to the ethical consideration of not withholding treatment, and given that our primary aim was to determine nerve regeneration and its relationship to functional recovery, our study used a prospective cohort rather than a trial design of randomising patients to surgical intervention or conservative management. Nevertheless, we continued to follow up thirteen patients who decided not to undergo surgery. The small sample and the absence of randomisation with lower age and symptom severity in non-operated patients precludes a direct comparison. However, the progressive structural decline of small nerve fibres that we identified in this group is however intriguing, and the potential implications for management of these patients warrants further investigation.

We have shown that PACAP is significantly upregulated following cutaneous reinnervation and can promote axon outgrowth in a human sensory neuronal model. Exploitation of PACAP as a treatment to promote neural regeneration *in vivo* will require consideration of optimal delivery given that systemic treatment can trigger migraine^41^. In fact, function blocking antibodies to PAC1 are undergoing clinical testing for migraine treatment^42^ and given our findings, care will need to be taken that such treatments do not impair sensory neuron function in the context of coexisting neuropathy. One means of optimising delivery to promote neurite outgrowth of small fibres would be to develop small molecule agonists of PAC1 which can be given locally as has been achieved for other growth promoting molecules such as GDNF^43^.

## Materials and Methods

### Study Design

This study used a prospective longitudinal cohort design paired with *in vitro* experiments to determine the cellular and molecular correlates of nerve regeneration in humans. Seventy-three patients with clinically^44^ and electrodiagnostically^45^ confirmed CTS were recruited from the surgical waiting lists at Oxford University Hospitals NHS Foundation Trust. Patients were excluded if electrodiagnostic testing revealed abnormalities other than CTS, if another medical condition affecting the upper limb and neck was present (e.g., hand osteoarthritis, cervical radiculopathy), if there was a history of significant trauma to the upper limb or neck, or if CTS was related to pregnancy or diabetes. Patients undergoing repeat carpal tunnel surgery were excluded from the study. After inclusion in this cohort, 13 patients opted out of surgery with the study team having no role in these patients’ decisions. While these patients were not the main focus of the study, we continued to follow them over time and report their data separately. A cohort of twenty healthy volunteers (proportionally age- and gender-matched to the CTS surgery group) without any systemic medical condition, or a history of hand, arm or neck symptoms, served as healthy controls. The required sample size was a priori determined with G-Power software^46^ for the primary outcome measure of small fibre regeneneration following surgery using previous data from the cross-sectional Oxford CTS cohort^9^. Fifty-nine patients were required to detect a 20% increase in IENFD with 80% power, significance set at α=0.05 and an effect size of 0.328.

The study was approved by the London Riverside national research ethics committee (Ref 10/H0706/35), and all participants gave informed written consent after the nature and possible consequences of the study has been explained. Whereas the healthy participants only attended a single session, all patients (including those that did and did not undergo surgery) attended two appointments; one at baseline and a follow up appointment (∼6 months after surgery/baseline appointment). In patients with CTS, the hand that was operated (surgery group) or the hand that was more affected (no-surgery group) was tested, whereas the non-dominant hand was tested in healthy controls. All clinical measurements were collected by the same experienced examiner to ensure consistency.

### Symptom and function questionnaires

Patients with CTS completed the Boston Carpal Tunnel Questionnaire (symptom and function scale; 0 = no symptoms/disability, 5 = very severe symptoms/disability)^47^ to determine symptom severity and functional deficits. Clinical recovery was determined with two global rating of change scales (GROC)^48^ evaluating changes in hand symptoms and function. The GROC scales range from ‘a very great deal better’ (+7), to ‘a very great deal worse’ (−7). A change of 5 or more points (a good deal better/worse) is considered a clinically meaningful change^49^. In addition, we calculated the difference in the Boston questionnaire subscales (follow-up minus baseline).

### Electrodiagnostic tests

Electrodiagnostic tests were performed with an ADVANCE system (Neurometrix, Waltham, MA, USA). Orthodromic sensory latencies and amplitudes were recorded over the digit to wrist segments for the median (index finger), ulnar (little finger) and superficial radial nerve (snuffbox) as previously described (*9*). Compound motor potentials (CMAP) were recorded over the abductor pollicis brevis and adductor digiti minimi by stimulating at the elbow as well as at the wrist. Radial CMAPs (extensor indicis proprius) were recorded by stimulating over the spiral groove. Electrodiagnostic testing was graded according to the scale by Bland^45^ as mild (2), moderate (3), severe (4), very severe (5) or extremely severe (6). To determine the presence of a very mild CTS (1), we included two sensitive tests: the presence of a ‘double peak’ during combined ulnar and median sensory stimulation at the ring finger and recording at the wrist^50^ and the presence of prolonged lumbrical to interossei motor latency difference >0.4ms when measured over a fixed distance of 8cm^51^. Hand temperature was standardised to >31 degrees Celsius. During data analysis, absent sensory and motor recordings were replaced with values of zero for amplitudes but excluded from analysis of latencies and nerve conduction velocities to prevent inflated results.

### Quantitative sensory testing (QST)

QST was performed according to the standardised protocol published by the German research network of neuropathic pain^52^. Cold and warm detection (CDT, WDT) and cold and heat pain thresholds (CPT, HPT) as well as thermal sensory limen (TSL) were measured with a thermotester (Somedic, Sweden). We also examined mechanical detection (MDT), vibration detection (VDT), mechanical pain thresholds (MPT), pressure pain thresholds (PPT), wind up ratio (WUR) and mechanical pain sensitivity (MPS). The number of paradoxical heat sensations during TSL and the presence of dynamic mechanical allodynia was quantified. All measurements apart from PPT and VDT were performed over the palmar aspect of the proximal phalanx of the index finger representing the affected median nerve territory. PPT was recorded over the thenar eminence and VDT over the palmar side of the distal end of the second metacarpal. To achieve normally distributed data, all QST parameters expect for CPT, HPT and VDT were log-transformed before statistical comparison^53^. A small constant of 0.1 was added to the mechanical pain sensitivity to avoid loss of zero-rating values. Most QST measures have good to excellent long-term reliability except for WUR, which has fair retest reliability over a period of 4 months^54, 55^.

### Histological analysis of target tissue

#### Skin biopsies

Two serial skin biopsies (3mm diameter) were taken 6 months apart on the ventrolateral aspect of the proximal phalanx of the index finger, innervated by the median nerve. The second biopsy was taken a few millimetres more proximal, avoiding the primary biopsy site. One surgical patient received Warfarin for unrelated reasons and no follow-up biopsy was thus taken. The biopsies were performed under sterile conditions following local anaesthesia with 1% lidocaine (1-1.8ml). Whereas half of the skin biopsy was immediately snap frozen for transcriptional analysis (see below), the other half was fixed in fresh periodate-lysine-paraformaldehyde for 30 minutes before being washed in 0.1M phosphate buffer and cryoprotected in 15% sucrose in 0.1M phosphate buffer. After embedding in OCT, the tissue was frozen and stored at -80 degrees. To assure consistency, each histological analysis was performed by the same blinded observer.

#### Intraepidermal nerve fibre density (IENFD)

Fifty μm thin sections were cut on a cryostat and stained using a previously described protocol^9^. Serial biopsies of the same patient were processed in the same batch to minimise variability. Sections were blocked with 5% fish gelatine before incubation with the primary antibodies for myelin basic protein (MBP, Abcam, 1:500) and protein gene product 9.5 (PGP, Ultraclone, 1:1000) overnight. Secondary antibodies were also incubated overnight (Alexa 488 1:1000, Abcam; Cy3 Stratech 1:500). The PGP antibody from Ultraclone was discontinued during our study. Benchmarking revealed that the PGP antibody from Zytomed (1:200) provided comparable staining quality and epidermal nerve fibre counts in biopsies of the same patients, and was therefore used for subsequent analyses.

Intraepidermal nerve fibre density (IENFD) was established by counting 3 sections per participant down the microscope, while strictly adhering to established counting criteria^56^. Intrarater reliability to determine IENFD was examined by processing and counting n=23 skin samples twice, 1 week apart. Intraclass correlation coefficient revealed almost perfect intra-rater agreement for IENFD (ICC 2,1 0.913 (95% confidence interval 0.791-0.963), p<0.0001).

#### Myelinated fibre integrity

We used several approaches to evaluate myelinated fibre integrity. First, we quantified the number of Meissner corpuscles per mm epidermis as previously reported^9^. Second, dermal nerve bundles containing myelinated fibres were expressed as a ratio of total PGP 9.5+ nerve bundles containing at least five PGP+ axons per mm^2^ dermis (excluding the subepidermal plexus). Third, we measured the length of nodes of Ranvier in the subepidermal plexus of MBP/PGP double-stained sections in up to 10 sections per participant. This was achieved with the Zen black software (Zeiss, UK) on confocal image stacks taken at 40x magnification as previously reported^9^. Nodes were classified as elongated if they exceeded 6.1μm in length^57^. If fewer than 15 nodes were identified, patients were excluded from the analysis (n=8).

#### Subepidermal plexus length

Estimation of nerve fibre length density in the subepidermal plexus was performed on PGP stained skin sections using an Olympus BX51 microscope (60× oil immersion lens) and newCAST stereological software (Visiopharm, Hoersholm, Denmark) in a blinded fashion. A virtual plane probe was used, as it generates isotropy of the test planes and allows length estimation of tubular objects in thick sections with arbitrary directions. The region of interest was the subepidermal plexus, here defined as an area deep to the epidermal-dermal junction and 200 µm towards the dermis. Papillae were excluded as they contain Meissner corpuscles, which are not part of the subepidermal plexus and their irregular distribution may have biased the results. Briefly, 3D sampling boxes are superimposed over the skin section, thereby generating randomized isotropic virtual planes in systematically sampled fields of view (for detailed methodological description see^58^). Sampling box height was set to 15 µm and the box area size to 7,908 µm^2^. Sampling steps were 85 × 70 µm (in the x and y-direction), with a plane separation distance of 25 µm. Four sections were typically counted per patient (ranging from 3-6).

### Statistical analysis of clinical and histological data

Data were analysed using SPSS (version 24, IBM) and R software^59^. Normal distribution of clinical and histological data was established by visual inspection and with the Kolmogorov-Smirnov test for normality.

The main statistical comparison concerned the pre-vs post-surgical data. This was achieved using two-sided paired t-tests or Wilcoxon signed rank tests for parametric and non-parametric data respectively. In order to establish differences at baseline and the extent of recovery, pre- and post-surgical data of patients with CTS were compared with healthy control data using two-sided independent t-tests or Mann Whitney U tests where appropriate. In order to control for false discovery rate (FDR) of multiple testing, Benjamini-Hochberg correction was applied for three comparisons with the false positive rate set to 5%^60^. Unadjusted p-values are reported for ease of interpretation. Data of non-operated patients were analysed and reported separately using the same statistical analyses as for operated patients.

### Correlation of histological findings with changes in symptomatology/function

Spearman’s correlation analyses were performed in the full patient cohort to determine associations between the extent of clinical recovery and neural regeneration. The extent of neural regeneration was determined by calculating the difference (follow-up minus baseline) of the histological parameters which significantly changed after surgery (IENFD, percentage of elongated nodes, mean nodal length). Clinical recovery was evaluated with the global rating of change questionnaire (for symptoms and function) as well as with the difference in Boston questionnaire (follow-up minus baseline for symptoms and function subscales). Benjamini-Hochberg correction was used to correct for multiple testing (12 correlations), with unadjusted p-values reported. To aid interpretation, Boston questionnaire data were multiplied by -1 so that a positive correlation always corresponds with more pronounced clinical recovery.

### Molecular analysis of target tissue

#### RNA sequencing

Half of the skin biopsies of 47 patients with CTS (n=29 females) before and after surgery were snap frozen in liquid nitrogen. RNA was extracted using a combination of phenol extraction and column purification. Samples were homogenized in Trizol and mixed with chloroform before spinning to induce phase separation. The aqueous liquid phase containing the nucleic acids was removed and added to the columns of the High Pure RNA tissue kit (Roche Diagnostics, UK). RNA was purified using repeated wash steps and DNAse treatment. The extracted RNA was sent for sequencing at the Wellcome Trust Centre for Human Genetics in Oxford. All samples were normalised to 650ng prior to library preparation using the TruSeq Stranded Total RNA Library Prep Kit (Illumina, UK) and poly-A enrichment. Paired-end sequencing was performed using the HiSeq4000 platform with a read length of 75bp.

RNA-sequencing data were mapped to the GRC.h.38 Human Genome using the STAR aligner^61^ with the ENCODE standard options. Read counts were calculated at the gene level using HTSeq^62^ and the ENSEMBL gene set annotation GRC.h.38.88. Raw counts were normalised using the effective library size, and for visualisations and associations with phenotypes they were transformed using the variance stabilising transformation (VST) in R using DESeq2^63^. Library size normalised gene counts were fitted to the negative binomial distribution and hypothesis testing was carried out using the Wald test. P-values were FDR corrected using the Benjamini-Hochberg procedure for multiple tests and the Independent Hypothesis Weighting (IHW)^64^. Moderated, i.e. shrunk towards zero, and non-moderated Log 2-fold changes (LFC) were used for hypothesis testing in differential expression (DE) analysis. We considered a gene as significantly DE if it had an adjusted p-value < 0.05 in at least two out of the three hypothesis testing procedures, i.e. moderated LFCs - FDR adjusted p-values, un-moderated LFCs - FDR adjusted p-values, un-moderated LFCs - IHW adjusted p-values.

Gene ontology enrichment for biological processes for DE genes was carried out in R using topGO^65^ and GSEA^66^. Hypothesis testing was performed using the weighted Fisher test and the significance cut-off was 0.01. The background gene list consisted of the 18068 genes expressed with > 0 counts in all samples.

#### Gene to regenerative phenotype association

Fold gene expression changes relative to baseline ((post-surgery – pre-surgery)/pre-surgery) and individual baseline expression of significantly DE genes were tested for association with fold changes and post versus pre differences in IENFD and nodal length. This was achieved by testing significant Pearson correlation coefficients and by binning the continuous outcomes into four categories corresponding to the four quarters as defined by the outcome quartiles with significance assessed using one-way ANOVA. In all associations of gene expression or fold change to phenotype, significance cut-off was an FDR adjusted p-value < 0.05. Raw and processed RNA-sequencing as well as basic phenotype data has been uploaded onto dbGAP (https://www.ncbi.nlm.nih.gov/projects/gap/cgi-bin/study.cgi?study_id=phs001796.v1.p1).

#### Droplet digital PCR to validate RNA sequencing

The findings of the RNA sequencing were validated in 78 paired samples pre and post surgery (n=39 patients) using digital droplet PCR (ddPCR). Complementary DNA was prepared using the Evoscript Universal cDNA Master kit (Roche Diagnostics, UK). TAQMAN assays (Thermo-Fisher, UK) for *ADCYAP1* (Hs00174950_m1) and *HPRT1* (Hs03929098_m1, reference) were run in duplicates with 1µl of target probe labelled with FAM added to 1µl of reference probe labelled with VIC in the same reaction using the ddPCR Supermix for Probes (Bio-Rad, UK). QuantaSoft v1.7.4.0917 software (Bio-Rad Laboratories) was used to extract data (copies/μl) and normalised gene expression was reported (ratios of target over reference data).

#### Localisation of PACAP in skin biopsies

The localisation of PACAP (encoded by *ADCYAP1*) expression in skin biopsies was evaluated using an immunofluorescent staining protocol adapted from Hannibal et al^67^. In brief, 50μm skin sections were mounted on glass slides and incubated in 10% goat serum in phosphate buffered saline and Triton-X for 30 minutes. Primary antibodies for PACAP (gift from Prof Jan Fahrenkrug, 1:5) and PGP9.5 (Zytomed, 1:200) were incubated overnight.

A biotinylated goat anti mouse antibody (Vector laboratories, 1:200) was then applied for 2 hours along with an Alexa Flour 546 anti rabbit antibody (Life technologies, 1:500). After five washes, the sections were incubated for 30 minutes with an avidin-biotin-horseradish peroxidase (VECTASTAIN® Elite ABC Kit, Vector laboratories) before washing four times. The sections were incubated for 10 minutes in FITC conjugated tyramide (Perkin Elmer, 1:100) diluted in 0.1M Borate buffer containing 0.0003% hydrogen perioxide. After 3 washes, slides were cover-slipped in mounting medium and imaged on a n Observer Z1 confocal imaging system (Zeiss).

### In-vitro expression and role of ADCYAP1/PACAP

#### RNA-sequencing of hiPSCd-sensory neurons

We interrogated our data from a previous RNA sequencing experiment^68^ in human fibroblast-derived induced pluripotent stem cell-derived sensory neurons which were generated as previously described^13^. Sequencing was performed using the Illumina HiSeq4000 paired-end protocol with 75bp reads. Reads were mapped on the Hg38 (GRCh38) human genome using STAR with the default ENCODE options. Gene counts were calculated for the GRCh38.88 gene set annotation using HTSeq. Data is available in the GEO series GSE107181. We evaluated *ADCYAP1* and its receptor *PAC1* expression levels compared to other genes.

#### Differentiation of human induced pluripotent stem cell (iPSC)-derived sensory neurons

The iPSC line NHDF is a control line derived from a healthy 44-year-old female^69^. Dermal fibroblasts from this individual were purchased from Lonza (CC-2511) which were then used for reprogramming to pluripotency. Lonza provide the following ethics statement: ‘These cells were isolated from donated human tissue after obtaining permission for their use in research applications by informed consent or legal authorization.’ The human iPSCs derived from these fibroblasts were generated as control lines for part of a larger-scale project (Ethics committee: NRES Committee South Central – Berkshire UK, REC 10/H0505/71). The fibroblasts were differentiated to sensory neurons as described previously^12, 13, 70^. In brief, cells were plated at high density following Versene EDTA (ThermoFisher) passaging. Neural induction was initiated in KSR medium (Knockout-DMEM, 15% knockout-serum replacement, 100 µM β-mercaptoethanol, 1% nonessential amino acids 1%, Glutamax (ThermoFisher)) by dual SMAD inhibition (SB431542 (Sigma, 10 µM) and LDN-193189 (Sigma, 100 nM). Three additional small molecules were introduced on day 3: CHIR99021 (Sigma, 3 µM), SU5402 (R&D Systems, 10 µM) and DAPT (Sigma, 10 µM) and dual SMAD inhibitors were withdrawn on day 5. KSR medium was gradually transitioned in 25% increments to neural medium (N2/B27-Neurobasal medium, 2% B27 supplement, 1% N2 supplement, 1% Glutamax, (ThermoFisher)) over an 11-day period. Cells were subsequently dissociated and replated onto glass coverslips in neural medium supplemented with growth factors at 25ng/ml (BDNF; ThermoFisher, NT3, NGF, GDNF; Peprotech). CHIR90221 was included for 4 further days, while SU5402 and DAPT were no longer included at this point. Phenol-free Matrigel (Corning, 1:300 dilution) was included from 25 days onward. Medium changes were performed twice weekly. Neurons were matured for 27±1 weeks before performing neurite outgrowth assays.

#### Human sensory neuron regeneration model

In order to determine a potential regenerative role of PACAP in human primary sensory neurons, we dissociated mature hiPSCd-sensory neurons and assessed their ability to regenerate neurites, as we have previously described^14^. Neurons were enzymatically treated for 30 min with 0.1% Trypsin (ThermoFisher), followed by mechanical dissociation with a fire polished glass pipette. Single cells were re-plated onto matrigel treated coverslips at low density in the presence of 10nM PACAP protein (AnaSpec) or 0.01% DMSO (Sigma), before fixation and analysis by immunocytochemistry 18 hours later.

#### Quantification of neurite outgrowth

One differentiation of mature hiPSCd-sensory neurons was used for all studies. Neurons were taken from individual wells of a 24-well plate and dissociated separately to form each experimental unit (*n*). Two dissociation experiments were performed. Average neurite outgrowth was calculated from at least two technical replicates (individual coverslips of replated neurons) per *n*. hiPSCd-sensory neurons treated with vehicle (*n*=7) and PACAP (*n*=6) were immunostained with NF200 using immunohistochemistry. Briefly, the NF200 antibody (Sigma, 1:500) was added to hiPSCd-sensory neurons overnight at room temperature. Neurons were then washed 3 times with PBS + TritonX and a secondary anti-mouse antibody Alexa Flour 488 (Life Technologies: 1:1000) was added for 2 hours at room temperature. The neurons were then washed 3x with PBS + TritonX and the coverslips were mounted with VectorShield on slides and sealed.

To determine neurite outgrowth, 20x magnified images of randomly selected individual hiPSCd-sensory neurons were taken on an Observer Z1 imaging system (Zeiss) and analysed for neurite length using WIS-Neuromath software^71^. Only neurons with a neurite at least half the diameter of the cell body were included. Neurite length was determined in μm per cell and the resultant neurite length from individual neurons were averaged per dissociation (4-18 neurons per dissociation). A Mann-Whitney U test was used to compare neurite length between PACAP treated or Vehicle treated groups.

The localization of PAC1 in hiPSCd-sensory neurons was examined with double labelling with NF200 (Abcam, 1:2000) and PAC1 (Cambridge Bioscience, 1:50) using the same immunohistochemistry protocol outlined above.

## Acknowledgments

We would like to thank all participants for volunteering in our study. The generous donation of the PACAP antibody by Prof Jan Fahrenkrug is gratefully acknowledged. This work was supported by the NIHR Biomedical Research Centre, Oxford. AB Schmid was funded by an advanced postdoc mobility Fellowship from the Swiss National Science Foundation (P00P3-158835) and received project funding from the early career research grant awarded by the International Association for the Study of Pain. DL Bennett is a Wellcome senior clinical scientist (ref. no. 095698z/11/z and 202747/Z/16/Z) and is supported by the Wellcome Trust Pain Consortium Strategic Award. O Sandy-Hindmarch is supported by a studentship from Immunocore Ltd. D Furniss was supported by an Intermediate Clinical Fellowship from the Wellcome Trust (097152/Z/11/Z). Disclaimer: The views expressed are those of the authors and not necessarily those of the NHS, the NIHR or the Department of Health.

## Author contributions

ABS designed the study, acquired funding, collected the data, performed and supervised analysis and wrote the manuscript. GB designed the bioinformatic workflows and analysed the RNA-sequencing data. KW and PK performed part of the histological analyses. OSH performed the ddPCR experiments and the PACAP staining in the skin. OSH, GW, LMD and AC performed the in vitro iPSC experiment. JB, MN, AW and DW recruited the patients. DLB designed the study, acquired funding and supervised the data analysis and manuscript preparation. All authors commented on the final version of the manuscript.

## Materials and Correspondence

Correspondence and requests for materials should be addressed to ABS or DLB.

## Competing interests

The authors declare no competing interests.

## Data availability

The data that support the findings of this study are deposited in the dbGaP repository (https://www.ncbi.nlm.nih.gov/projects/gap/cgi-bin/study.cgi?study_id=phs001796.v1.p1).

## Code availability

Standard scripts using already published R packages were used and cited throughout the manuscript.

## Notes

#### Summary of Updates

change of wrong OrchID for co-author McDermott

https://www.ncbi.nlm.nih.gov/projects/gap/cgi-bin/study.cgi?study_id=phs001796.v1.p1

