## Supplementary material for "Molecular and cellular correlates of human nerve regeneration: *ADCYAP1* encoding PACAP enhances sensory neuron outgrowth": Table S1

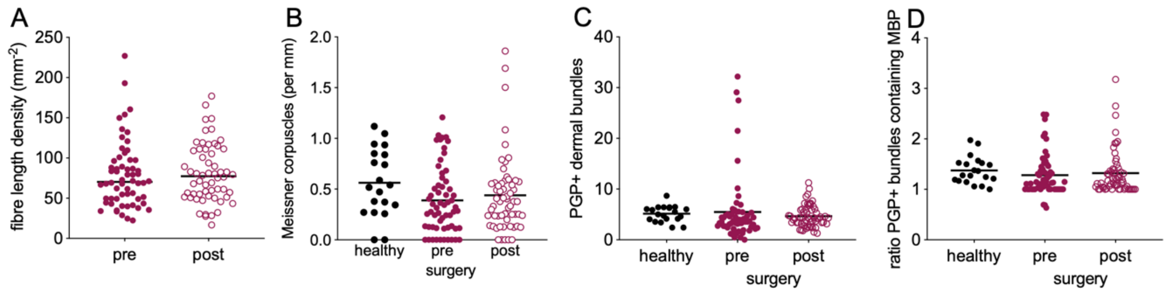

**Fig S1:** Dermal innervation is comparable between healthy controls and CTS patients and does not change after surgery. **(A)** Subepidermal plexus fibre length density **(B)** Meissner corpuscle density **(C)** Protein gene product 9.5 (PGP)+ dermal bundles and **(D)** ratio of PGP+ dermal bundles containing myelin basic protein (MBP).

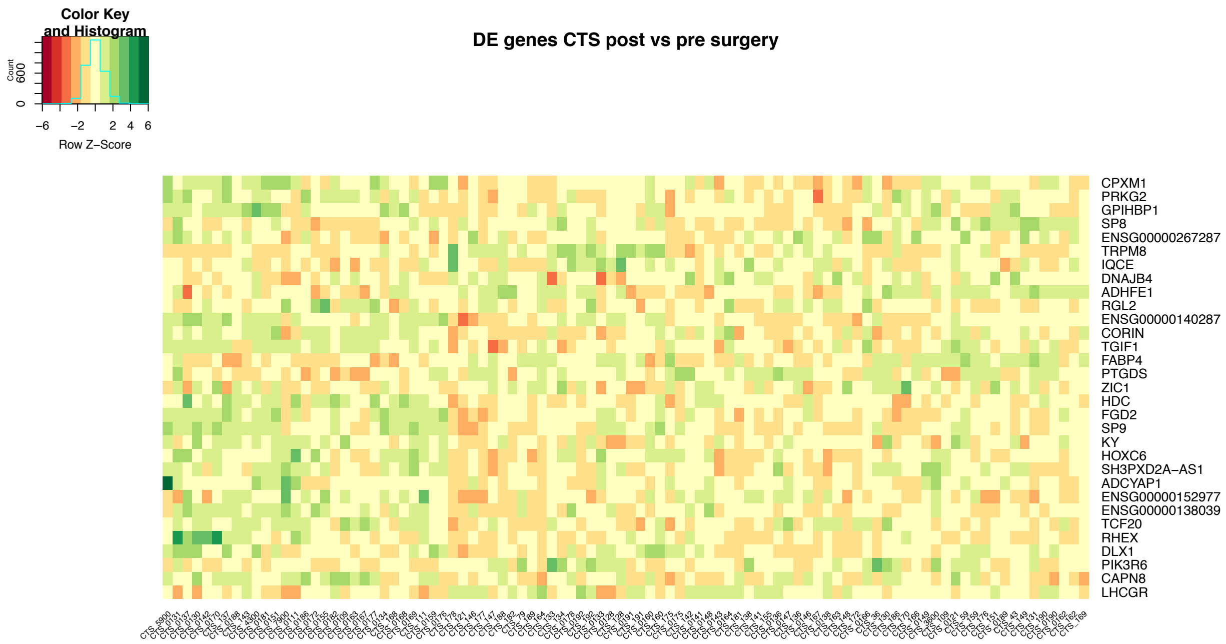

**Fig S2:** Heatmap of differentially expressed genes identified in the RNA sequencing experiment in human skin (n=47). Heatmap of the relative expression changes based on centered and scaled regularised log2 transformed gene counts. Color key shows the mapping between colours and z-score transformed gene expression.

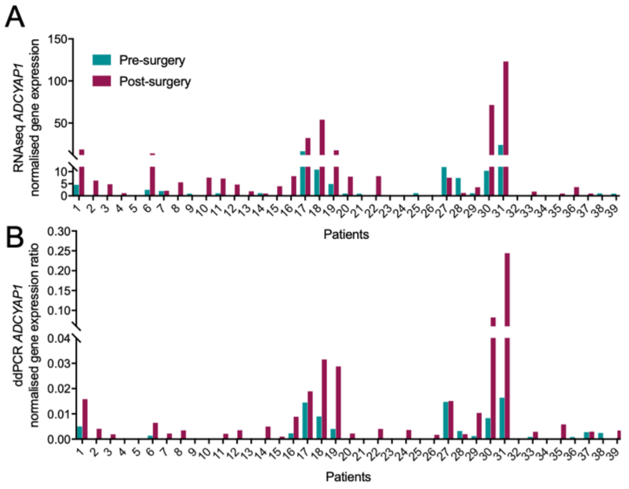

**Fig S3:** Validation of *ADCYAP1* mRNA expression using droplet digital PCR. **(A)** Normalised gene expression levels of the RNA sequencing align with **(B)** normalised gene expression ratio of the droplet digital PCR experiment (n=39).

Table S1: Quantitative sensory testing data of patients pre- and 6 months post-surgery as well as healthy controls.

| | surgery group | | | healthy controls | | |
|---|---|---|---|---|---|---|
| | pre | post | p-value (pre post) | baseline | p-value (HC-pre) | p-value (HC-post) |
| CDT (°C) | -5.66 (4.95) | -3.38 (2.33) | **<0.0001** | -2.28 (0.97) | **<0.0001** | **0.033** |
| WDT (°C) | 5.25 (3.89) | 4.40 (3.24) | 0.120 | 3.01 (1.66) | **<0.0001** | **0.022** |
| TSL (°C) | 11.15 (8.38) | 8.74 (5.73) | **0.002** | 5.53 (2.89) | **0.002** | **0.023** |
| MDT (mN) | 8.45 (19.00) | 3.01 (8.90) | **<0.0001** | 0.38 (0.26) | **<0.0001** | **0.006** |
| VDT (x/8) | 7.45 (0.74) | 7.78 (0.54) | **<0.0001** | 7.88 (0.32) | **0.015** | 0.429 |
| CPT (°C) | 9.42 (7.00) | 11.44 (7.01) | **0.046** | 7.88 (0.25) | 0.484 | 0.698 |
| HPT (°C) | 44.06 (3.94) | 43.74 (3.45) | 0.481 | 10.72 (7.56) | 0.484 | 0.664 |
| MPT (mN) | 180.4 (125.7) | 157.2 (103.3) | 0.264 | 153.44 (85.33) | 0.657 | 0.357 |
| MPS (0-100) | 0.59 (1.11) | 0.41 (0.39) | 0.569 | 0.56 (1.00) | 0.782 | 0.386 |
| PPT (kPa) | 180.4 (125.7) | 157.3 (103.3) | 0.221 | 351.36 (86.22) | 0.747 | 0.452 |
| WUR (ratio) | 2.53 (2.70) | 2.53 (2.24) | 0.810 | 2.04 (1.21) | 0.402 | 0.302 |

CDT, cold detection threshold; WDT, warm detection threshold; TSL, thermal sensory limen; MDT, mechanical detection threshold; VDT, vibration detection threshold; CPT, cold pain threshold; HPT, heat pain threshold; MPT, mechanical pain threshold; MPS, mechanical pain sensitivity; WUR, windup ratio; PPT, pressure pain threshold. Data are presented as mean (standard deviation) for untransformed data (CPT, HPT, VDT) and retransformed mean for log-tranformed data. P-values reflect statistics done on log transformed data as appropriate comparing pre and postoperative data (pre post), healthy controls with CTS patients presurgery (HC-pre) and healthy controls with CTS patients postsurgery (HC-post). P-values that were significant following Benjamini-Hochberg correction are highlighted in bold.

Table S2: Clinical, quantitative sensory testing and histological outcomes in CTS patients who did not undergo an operation.

| | baseline | follow-up | p-value |
|---|---|---|---|
| **Questionnaires** | | | |
| Mean VAS (SD) | | | |
| Pain | 2.3 (3.0) | 1.1 (1.5) | 0.268 |
| Numbness | 3.8 (3.2) | 1.6 (2.2) | 0.063 |
| Paraesthesia | 3.9 (2.8) | 2.6 (3.0) | 0.271 |
| Mean Boston scale (SD) | | | |
| Symptoms | 2.3 (0.6) | 2.0 (0.6) | 0.215 |
| Function | 1.9 (0.6) | 1.6 (0.5) | 0.171 |
| Mean NPSI (SD) | 9.7 (7.7) | 5.6 (5.8) | 0.186 |
| **Quantitative sensory testing** | | | |
| CDT (°C) | -3.40 (1.83) | -3.23 (1.65) | 0.805 |
| WDT (°C) | 4.07 (2.39) | 3.96 (1.98) | 0.914 |
| TSL (°C) | 6.53 (2.72) | 6.90 (3.73) | 0.874 |
| MDT (mN) | 1.27 (1.48) | 1.10 (1.16) | 0.980 |
| VDT (x/8) | 7.44 (0.63) | 7.28 (1.15) | 0.443 |
| CPT (°C) | 15.31 (8.06) | 11.38 (8.68) | **0.022** |
| HPT (°C) | 41.12 (2.19) | 42.54 (3.00) | 0.135 |
| MPT (mN) | 100.5 (66.9) | 111.4 (92.7) | 0.565 |
| MPS (0-100) | 0.95 (0.84) | 0.56 (0.59) | **0.042** |
| PPT (kPa) | 325.5 (97.3) | 356.4 (109.7) | 0.069 |
| WUR (ratio) | 1.64 (0.66) | 2.24 (0.81) | **0.033** |
| **Histological parameters** | | | |
| IENFD | 6.94 (3.09) | 5.61 (3.41) | **0.021** |
| Meissner corpuscle density | 0.47 (0.31) | 0.20 (0.24) | **0.010** |
| Subepidermal plexus nerve fibre length | 137.0 (176.5) | 132.6 (110.1) | 0.433 |
| PGP+ dermal axon bundles | 5.74 (4.20) | 4.42 (1.98) | 0.299 |
| Ratio of PGP+ dermal bundles containing MBP | 1.45 (0.42) | 1.57 (0.60) | 0.710 |
| Nodal length (median [IQR]) | 2.00 [1.01] | 2.72 [2.63] | **0.023** |
| % elongated nodes (median [IQR]) | 0.00 [4.55] | 6.66 [22.22] | 0.063 |
| Internodal length | 82.8 (25.88) | 70.24 (16.83) | 0.062 |
| G-ratio | 0.84 (0.07) | 0.89 (0.06) | 0.107 |

CDT, cold detection threshold; WDT, warm detection threshold; TSL, thermal sensory limen; MDT, mechanical detection threshold; VDT, vibration detection threshold; CPT, cold pain threshold; HPT, heat pain threshold; MPT, mechanical pain threshold; MPS, mechanical pain sensitivity; WUR, windup ratio; PPT, pressure pain threshold; IENFD: intraepidermal nerve fibre density (per mm epidermis; PGP: protein gene product; MBP: myelin basic protein. Data are presented as mean (standard deviation) unless indicated otherwise.

Table S3: Differentially expressed genes in skin post compared to pre surgery

| GeneID | baseMean | log2FoldChange | padjBH | padjW | padjBetaPrior | Symbol |
|---|---|---|---|---|---|---|
| ENSG00000141433 | 8.29462916 | 1.87901203 | 0.00696535 | 0.00641312 | 0.00010901 | ADCYAP1 |
| ENSG00000138039 | 4.65862691 | 1.25438865 | 0.01145755 | 1 | 0.01401089 | LHCGR |
| ENSG00000152977 | 4.89145385 | 1.11254793 | 0.00842065 | 0.01940691 | 0.03253888 | ZIC1 |
| ENSG00000088882 | 92.5184023 | 0.92190964 | 0.00103854 | 0.00064558 | 7.93E-06 | CPXM1 |
| ENSG00000280113 | 8.0167541 | 0.85235048 | 0.00848225 | 0.02412323 | 0.02563488 | NA |
| ENSG00000140287 | 203.616122 | 0.72241168 | 0.00983556 | 0.0074644 | 0.03253888 | HDC |
| ENSG00000267453 | 19.7973425 | 0.67273101 | 0.02111925 | 0.01940691 | 0.02522041 | LINC01835 |
| ENSG00000263961 | 44.8376836 | 0.62915302 | 0.00696535 | 0.00393004 | 0.01854591 | RHEX |
| ENSG00000170323 | 77.6103507 | 0.62247621 | 0.05612312 | 0.02862547 | 0.02855876 | FABP4 |
| ENSG00000276231 | 35.1351758 | 0.49145976 | 0.03282225 | 0.01705774 | 0.0435133 | PIK3R6 |
| ENSG00000277494 | 67.4781354 | 0.44054154 | 0.00848225 | 0.00518474 | 0.00504287 | GPIHBP1 |
| ENSG00000107317 | 135.740333 | 0.42046293 | 0.00762641 | 0.00421831 | 0.00796676 | PTGDS |
| ENSG00000138669 | 39.7099245 | 0.41788473 | 0.00328312 | 0.00174962 | 0.00376299 | PRKG2 |
| ENSG00000270605 | 48.9272349 | 0.41678525 | 0.03447244 | 0.13317736 | 0.02855876 | NA |
| ENSG00000145244 | 143.132317 | 0.30856247 | 0.03863323 | 0.01705774 | 0.03104537 | CORIN |
| ENSG00000147576 | 120.447618 | 0.30289174 | 0.00696535 | 0.00275544 | 0.00645172 | ADHFE1 |
| ENSG00000197757 | 287.995304 | 0.19212071 | 0.01723099 | 0.00975572 | 0.01708509 | HOXC6 |
| ENSG00000162616 | 506.036925 | 0.17310075 | 0.0373841 | 0.07851918 | 0.03208498 | DNAJB4 |
| ENSG00000177426 | 489.021911 | 0.13250093 | 0.01723099 | 0.02115744 | 0.01883526 | TGIF1 |
| ENSG00000237441 | 2611.98287 | 0.10001312 | 0.03596432 | 0.01705774 | 0.03104537 | RGL2 |
| ENSG00000144481 | 17.0057572 | -0.7848364 | 0.00762641 | 0.01705774 | 0.00504287 | TRPM8 |
| ENSG00000164651 | 220.844691 | -0.5629928 | 2.77E-09 | 7.80E-10 | 9.96E-09 | SP8 |
| ENSG00000203697 | 173.954774 | -0.4487044 | 0.05612312 | 0.02862547 | 0.02855876 | CAPN8 |
| ENSG00000217236 | 110.859221 | -0.3937448 | 0.01145755 | 0.00704249 | 0.02110566 | SP9 |
| ENSG00000267287 | 80.1845927 | -0.3506917 | 0.03354355 | 0.01705774 | 0.03423147 | NA |
| ENSG00000144355 | 489.75608 | -0.3277165 | 0.00049206 | 0.0010581 | 0.00020125 | DLX1 |
| ENSG00000174611 | 165.305979 | -0.2733539 | 0.02060243 | 0.01705774 | 0.0241341 | KY |
| ENSG00000146192 | 551.866588 | -0.2174852 | 0.03596432 | 0.0496477 | 0.03953373 | FGD2 |
| ENSG00000280693 | 188.838376 | -0.2155644 | 0.02876924 | 0.01940691 | 0.02855876 | SH3PXD2A-AS1 |
| ENSG00000100207 | 1378.70972 | -0.1335073 | 0.02341961 | 0.0366254 | 0.02495219 | TCF20 |
| ENSG00000106012 | 714.887208 | -0.0974793 | 0.00593115 | 0.00491958 | 0.00497218 | IQCE |

Table S4: Gene ontology (GO) terms for biological processes identified in skin

| GO.ID | Term | Annotated | Significant | Expected | weightFisher |
|---|---|---|---|---|---|
| GO:0045766 | positive regulation of angiogenesis | 117 | 3 | 0.2 | 0.00094 |
| GO:0009954 | proximal/distal pattern formation | 28 | 2 | 0.05 | 0.00099 |
| GO:0050999 | regulation of nitric-oxide synthase activity | 35 | 2 | 0.06 | 0.00154 |
| GO:0032611 | interleukin-1 beta production | 49 | 2 | 0.08 | 0.003 |
| GO:0042632 | cholesterol homeostasis | 52 | 2 | 0.09 | 0.00338 |
| GO:0006357 | regulation of transcription from RNA polymerase II | 1626 | 7 | 2.72 | 0.00632 |
| GO:0006810 | transport | 4071 | 9 | 6.82 | 0.00847 |
| GO:0042089 | cytokine biosynthetic process | 80 | 2 | 0.13 | 0.00961 |
| GO:0048706 | embryonic skeletal system development | 99 | 2 | 0.17 | 0.01177 |
| GO:0006821 | chloride transport | 69 | 2 | 0.12 | 0.01278 |
| GO:0001990 | regulation of systemic arterial blood pressure | 30 | 2 | 0.05 | 0.0159 |
| GO:0030326 | embryonic limb morphogenesis | 116 | 2 | 0.19 | 0.01591 |
| GO:0010623 | programmed cell death involved in cell development | 10 | 1 | 0.02 | 0.01663 |
| GO:0002003 | angiotensin maturation | 10 | 1 | 0.02 | 0.01663 |
| GO:0071285 | cellular response to lithium ion | 10 | 1 | 0.02 | 0.01663 |
| GO:0042368 | vitamin D biosynthetic process | 10 | 1 | 0.02 | 0.01663 |
| GO:0001660 | fever generation | 10 | 1 | 0.02 | 0.01663 |
| GO:0042033 | chemokine biosynthetic process | 10 | 1 | 0.02 | 0.01663 |
| GO:0045187 | regulation of circadian sleep/wake cycle | 10 | 1 | 0.02 | 0.01663 |
| GO:0030656 | regulation of vitamin metabolic process | 10 | 1 | 0.02 | 0.01663 |
